# Next Generation Aging Clock: A Novel Approach to Decoding Human Aging Through Over 3000 Cellular Pathways

**DOI:** 10.1101/2024.06.18.599632

**Authors:** Jianghui Xiong

## Abstract

This paper introduces ‘Next Generation Aging Clock Models,’ a new approach aimed at improving disease prediction by defining aging clocks for specific cellular components or pathways, rather than giving a single value for the entire human body. The methodology consists of two stages: a pre-training stage that creates 3,028 generic pathway aging models by integrating genome-wide DNA methylation data with gene ontology and pathway databases, and a fine-tuning stage that produces 30,280 disease-specific pathway aging models using DNA methylation profiles from 3,263 samples across 10 age-related diseases. Our findings show the model’s predictive power for various diseases. For example, the aging index of blood vessel endothelial cell migration can predict Atherosclerosis with an odds ratio of 80. Alzheimer’s disease can be predicted by the aging index of response to DNA damage stimulus, Major Depressive Disorder by the organization of the mitochondrion, breast cancer by DNA repair, and the severity of COVID-19 by neutrophil degranulation, with an odds ratio of 8.5. Additionally, a global analysis revealed that aging-related diseases can be categorized into nucleus aging (such as Alzheimer’s disease) and cytoplasm aging (such as Parkinson’s disease). This model provides a comprehensive view of aging from the organelle to the organ level using just a blood or saliva sample. This innovative approach is expected to be a valuable tool for research into aging-related diseases and for personalized aging interventions.

## Introduction

The concept of biological age, distinct from chronological age, has become significant in aging research. This emphasizes the importance of aging biomarkers in studying age-related diseases and implementing longevity interventions [1, 2]. Among the various models of biological aging, the DNA methylation clock is the most studied. This model predicts age based on the methylation status of specific CpG sites in the genome [3]. The discrepancy between predicted age and chronological age, referred to as ‘age acceleration’, is employed to explore disease correlations and predict health outcomes [4]. The first type of clock fits chronological age such as Horvath clock [5] and Hannum clock[6]. The second type of clocks estimate mortality risk such as GrimAge clock [7] and PhenoAge clock [8]. The third type of clock analyzes longitudinal data, such as DunedinPoAm [9] and Pace of Aging [10].

The traditional aging clock, which provides a single value for the entire human body and predicts disease and mortality risk, has limited disease prediction capabilities. Recently, an organ-specific clock that provides aging measures for ten organs has been introduced [11]. This multi-dimensional aging clock can forecast diseases specific to certain organs. For example, it has shown that accelerated heart aging boosts the risk of heart failure by 250%. A recent paper disclosed that the biological age of one organ could selectively impact the aging of other organ systems, thereby revealing a multiorgan aging network [12]. Additionally, a new causality-enriched epigenetic age was introduced, which can identify DNA methylation sites causally linked to aging-related traits, such as damage and adaptation [13].

Systems biology views the human body as a multi-level complex network, spanning from genes and cells to organs and systems. Given this, the basic biological process of aging leaves traces and records at each of these levels. With this in mind, we suggest expanding the concept of the biological aging clock to introduce the Pathway Aging Model. This model establishes an independent clock for each pathway, allowing for individualized assessment of aging status in each pathway. Given the thousands of pathways, we suggest that these advanced dimensional aging clocks could offer a more profound, detailed insight into aging and its relation to various illnesses. For complex conditions like neuropsychiatric diseases that impact multiple organs, the pathway clock could be especially beneficial. It can aid in understanding disease variation, facilitating disease subtyping and classification, and guiding personalized prevention and treatment strategies.

## Results

### The Framework of the Multi-Scale Models for Decoding Human Aging

The entire computational framework, as shown in Figure 1, consists of three stages: pre-training, fine-tuning, and application. In the pre-training stage, we aim to quantify the aging of all biological processes or structures theoretically. To start, we collected various gene sets reflecting biological functions. Gene ontology is a crucial source that defines biological processes, molecular functions, and cellular components. Additionally, there are numerous databases about signaling pathways, such as the Reactome pathway. This resulted in 9,509 candidate gene set definitions. For simplicity, we refer to all these gene sets as pathways. Ideally, we will define and calculate aging indexes for all gene sets.

**Figure 1.**
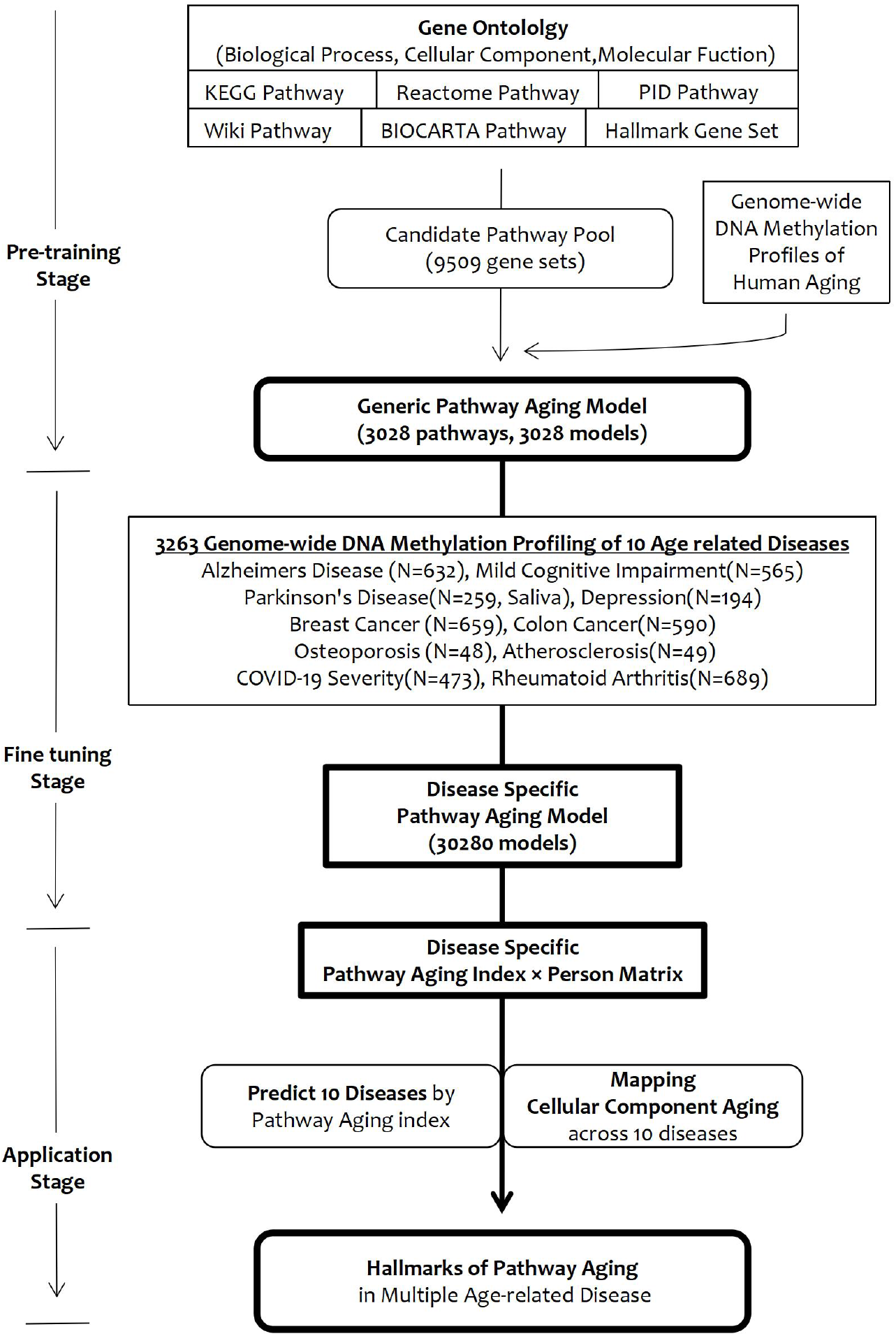
Workflow of the Next-Generation Aging Clock Model for Calculating Pathway Aging and Its Application in Predicting Age-Related Diseases.

We filtered the previously mentioned pathways, keeping only those that intersect with age-associated gene sets. This process retained 3,028 pathways, which include 1,295 Gene Ontology Biological Process (GOBP), 192 Gene Ontology Molecular Function (GOMF), 175 Gene Ontology Cellular Component (GOCC), and 184 Reactome signaling pathways, among others.

Each biological process or subcellular structure has genes that increase or decrease with aging. In relation to DNA methylation data, this usually means that the methylation of some gene promoter regions increases, while others decrease. Consequently, we used a whole genome DNA methylation dataset with age information to calculate the correlation between each gene and age. This resulted in age positively correlated genes, referred to as pos genes, and negatively correlated genes, known as neg genes.

Our main assumption is that as aging occurs, the difference between positive and negative genes in a biological process or structure increases. To measure this, we use a T-test to statistically evaluate the difference in methylation levels between positive and negative gene groups. A higher absolute t-score indicates a greater degree of aging. Thus, we use the t-score from the T-test to quantify the aging degree of a pathway. For each pathway, the aging index is represented by this t-score.

During the fine-tuning stage, we adjust the positive and negative genes for each pathway. We have used genome-wide DNA methylation data from 3,263 samples across 10 age-related diseases. For each disease, we identify genes significantly related to it at the gene level. We then intersect the positive gene list with disease-related genes to identify disease-specific positive genes (dsPos-genes) and similarly obtain disease-specific negative genes (dsNeg-genes). We use a T-test to statistically assess the difference in methylation levels between dsPos and dsNeg gene groups. This t-score is the disease-specific pathway aging index.

In this way, for each of the 3,028 pathways, we create 10 disease-specific models, resulting in a total of 30,280 disease-specific pathway aging models.

In the application stage, we use the specific model of each pathway-disease combination to transform the methylation profile of each disease into a new matrix, the PA-index×samples matrix, where each row represents a pathway aging index. We utilize these matrices to identify pathways that can predict diseases and to study the common characteristics of various diseases in terms of pathway aging.

### Pathway Aging as Predictive Factors of Disease

For 10 types of age-related diseases, we have obtained the list of pathways with the strongest predictive power, see supplementary file 1.

Figures 2 and 3 demonstrate how the pathway aging index can predict age-related diseases. For instance, Figure 2A shows two representative pathways predictive for Atherosclerosis. The orange box plot highlights a significant difference in the pathway aging index between the disease and control groups. The blue box plot shows the significant difference in the pathway aging index between different age groups (old vs. young). The data used is from GEO data GSE40279, as mentioned in the methods section. This data includes genome-wide DNA methylation profiling from whole blood samples of 656 individuals [6]. We use the age of 40 as a cutoff to divide the population into older and younger groups, comparing the significant difference in each pathway aging index.

**Figure 2.**
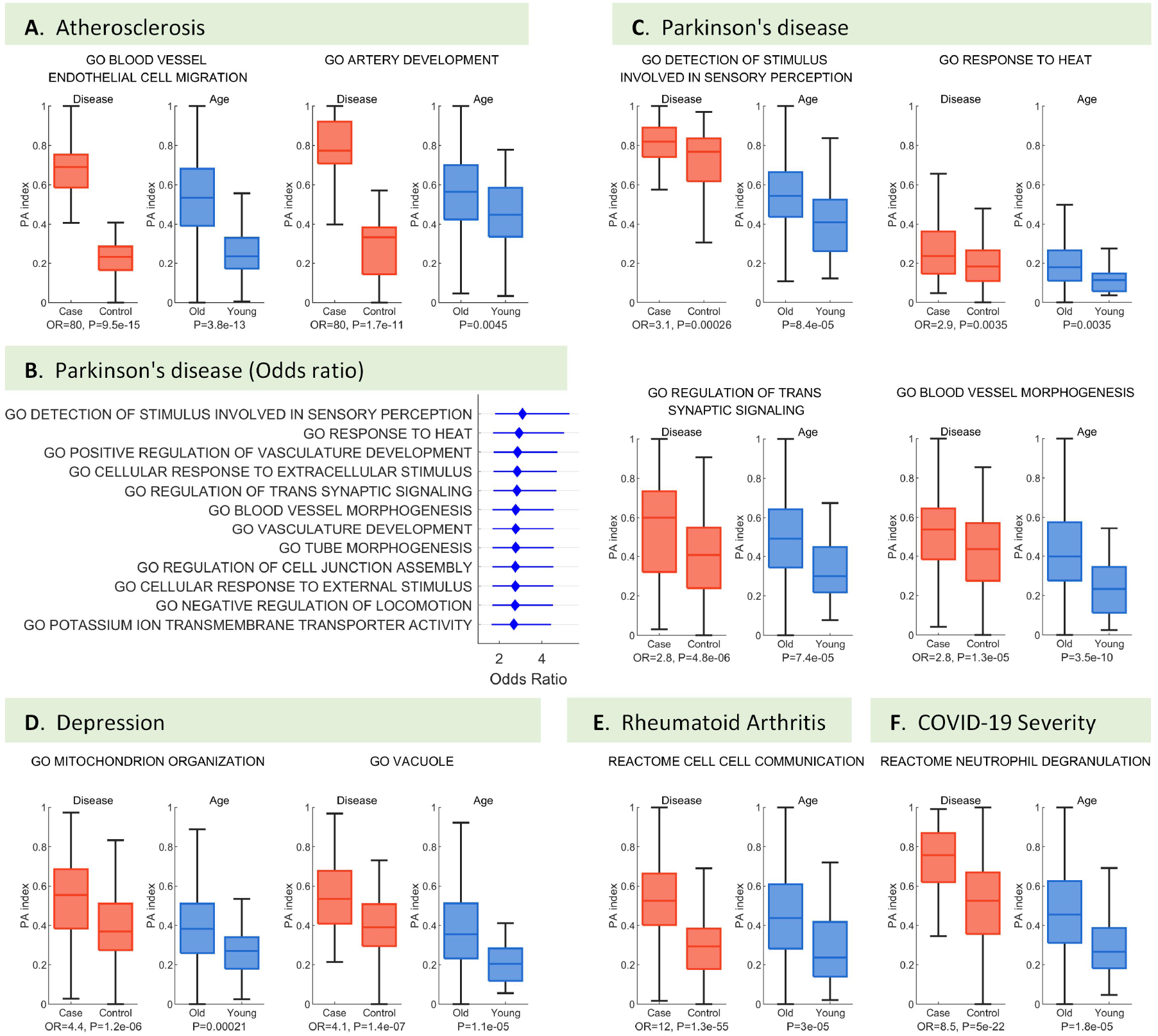
Pathway Aging as Predictive Factors of Disease. Subplots A-F illustrate key pathways that can predict various diseases. A. Atherosclerosis. B and C. Parkinson’s disease. D. Major depressive disorder. E. Rheumatoid arthritis. F. COVID-19 severity.

**Figure 3.**
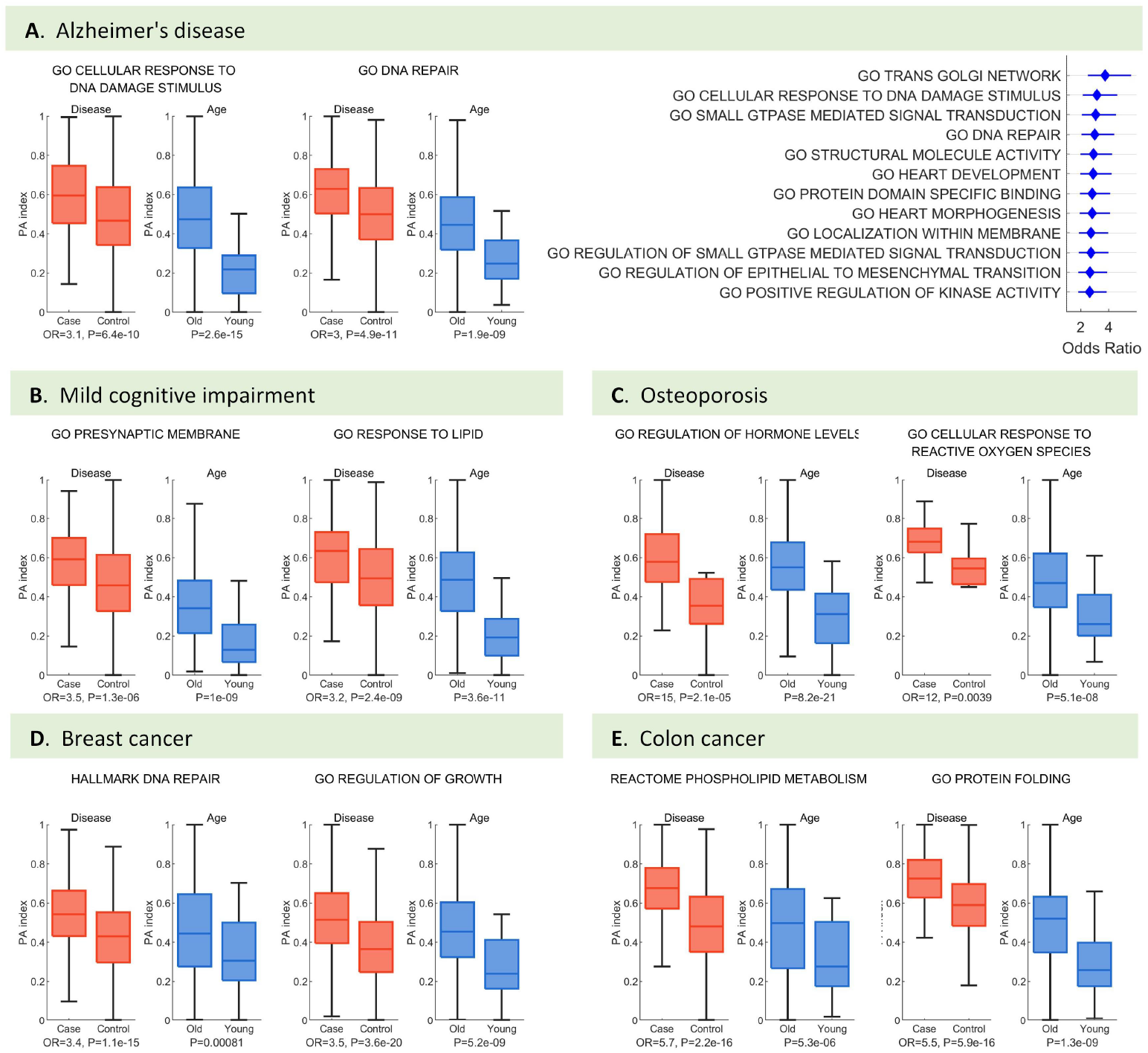
Pathway Aging as Predictive Factors of Disease (Part 2). Subplots A-E depict the representative pathways that can predict various diseases. A. Alzheimer’s Disease. B. Mild Cognitive Impairment. C. Osteoporosis. D. Breast Cancer. E. Colon Cancer.

The figure’s lower section includes the odds ratio (OR) and the p-value. The method to calculate the odds ratio is provided in the methods section. Briefly, it involves dividing a certain aging index into high and low groups based on its mean value. Then, it measures how frequently the disease appears in the high group compared to the low group. If the OR is greater than 1, a larger value indicates a higher likelihood of this variable becoming a risk factor for the disease, and the stronger its predictive power. The p-value signifies the significance of the difference between the disease and control groups’ aging index. It is obtained using a t-test.

As shown in Figure 2A, the pathway aging index of the Gene Ontology biological process term “blood vessel endothelial cell migration” can predict Atherosclerosis with an odds ratio of 80 and a p-value of 9.5e-15. The aging index of this pathway is also significantly different in aging groups (old vs. young), with a t-test p-value of 3.8E-13. Another variable with strong predictive power is GO artery development.

Endothelial cells in blood vessels play a significant regulatory role in Atherosclerosis, undergoing epigenetic reprogramming throughout the disease process [14]. Our findings suggest that a whole blood DNA methylation analysis can capture this crucial variable in Atherosclerosis progression.

Parkinson’s disease is a progressive disorder marked by motor dysfunction. Our analysis identified pathways with the highest odds ratios, shown in Fig. 2B. These include various Gene Ontology biological processes: detection of stimulus involved in sensory perception (OR=3.1), response to heat (OR=2.9), regulation of trans-synaptic signaling (OR=2.8), blood vessel morphogenesis (OR=2.8), and negative regulation of locomotion (OR=2.7). The pathway aging index for these pathways is also significantly higher in older individuals compared to younger ones (Fig. 2C). This suggests that these biological processes might support normal motor function, and their decline might contribute to the development of Parkinson’s disease. For instance, recent evidence shows that the body’s response to heat is relevant to Parkinson’s disease. A cross-sectional study with 247 Parkinson’s disease patients found that 78.9% reported increased heat sensitivity. Heat sensitivity in people with Parkinson’s disease significantly worsens both motor and non-motor symptoms, affecting daily activities, work effectiveness, and quality of life [15].

Beyond Gene Ontology biological processes, the aging index of GO cellular components is also a notable predictor for age-related diseases. For instance, in major depressive disorders, the aging index of mitochondrion organization (OR = 4.4) and vacuole (OR = 4.1) can predict these disorders (Fig. 2D). The link between mitochondrial dysfunction and depression is gaining more attention. For example, new insights into depression connect symptoms like fatigue and lack of motivation to changes in mitochondrial structure and function. This indicates that improving mitochondrial functions could be a promising approach for depression therapy [16].

The Reactome pathway also aids in predictive pathway definitions. For example, cell-cell communication defined by the Reactome pathway can predict Rheumatoid Arthritis with an odds ratio (OR) of 12 (Fig 2E). This is significant because cell-cell communications are crucial in rheumatoid arthritis (RA). Activation of the endothelium and immune cells initiates RA, leading to trans-endothelial cell migration, synovial fibroblast proliferation, and joint destruction [17].

Furthermore, neutrophil degranulation, as defined by the Reactome pathway, can predict COVID-19 severity with an odds ratio (OR) of 8.5 (Fig 2F). This process is significantly linked to COVID-19 severity as it contributes to the formation of neutrophil extracellular traps (NETs), which worsen immunothrombosis [18].

The correlation between other diseases and the pathway aging index is shown in Figure 3. Notably, Alzheimer’s disease and breast cancer both relate to the aging index of DNA damage repair related pathways. For example, Alzheimer’s disease could be predicted by the aging index of GO cellular response to DNA damage stimulus (OR=3.1) and GO DNA repair (OR=3.0, Fig 3A), while breast cancer could be predicted by the Hallmark gene set: DNA repair (OR=3.4, Fig 3D). This suggests that a decline in DNA repair is a common feature of age-related diseases. Genomic instability is a key hallmark of aging [19]. Targeting DNA damage could lead to unified interventions against age-related dysfunction and disease [20].

Similar to genomic instability, the loss of proteostasis is a significant hallmark of aging [19]. Our results indicate that the protein folding pathway aging index can predict colon cancer with an odds ratio of 5.5 (Fig. 3E). This finding is relevant because numerous studies demonstrate that the unfolded protein response (UPR) is closely linked to the occurrence and progression of colorectal cancer [21].

Another type of aging-related molecules is lipids. Lipids are essential for maintaining cellular balance and are becoming recognized as key regulators of aging. Disrupted lipid metabolism is linked to all known hallmarks of aging. For instance, ELOVL fatty acid elongase 2 (Elovl2), a gene with age-related epigenetic changes, influences aging by regulating lipid metabolism [22]. Our study also identified two significant lipid-related pathways. The GO response to the lipid aging index could predict mild cognitive impairment with an odds ratio of 3.2 (Fig. 3B), and the Reactome phospholipid metabolism aging index could predict colon cancer with an odds ratio of 5.7 (Fig. 3E).

Beyond molecular events, organ-level pathways also show potential for predicting age-related diseases. For example, our findings indicate that two heart related pathways, heart development and heart morphogenesis, may serve as predictors for Alzheimer’s disease (Fig 3A, right panel, Line 6 and Line 8). These results support the heart-brain axis theory, which connects the cardiovascular and cerebrovascular systems and influences both cardiovascular disease and Alzheimer’s [23]. This connection suggests that heart issues could lead to cognitive decline, including Alzheimer’s.

In addition to interactions between organs, broader regulation is managed through systemic circuits governed by hormones that link various organs across the body.We also found that the GO regulation of hormone levels aging index could predict osteoporosis (odds ratio 15, Fig. 3C).

These findings suggest that DNA methylation detection in whole blood can generally reflect pathway aging and is linked to diseases in various types of organs and tissues. Interestingly, the Parkinson’s disease analysis is based on saliva samples, implying that saliva, like whole blood, could also be a valuable source of information for decoding whole body pathway aging.

### Statistical Analysis of Gene Ontology Categories in Aging-Related Diseases

The Gene Ontology knowledgebase is a comprehensive, up-to-date computational model of biological systems, spanning from molecular pathways to cellular and organismal levels. It is well-structured and computer-readable, offering a detailed representation of current scientific knowledge about the structure and function of the human body. Defining an aging index for each Gene Ontology term is very useful for generating a multi-scale view of aging-related diseases.

We aim to identify the categories or dimensions of gene ontology that are most relevant to aging and the mechanisms of age-related diseases. Gene ontology categories include three main areas: Biological Process (GOBP), which outlines the pathways and larger processes to which these gene products contribute, Molecular Function (GOMF), which details the molecular activities of individual gene products and Cellular Component (GOCC), which identifies where these gene products are active. In the signaling pathway database, the Reactome pathway defines the most pathways, and we have included it in this analysis.

For each disease, we ranked pathways based on the odds ratio of the pathway aging index in predicting the disease. We then identified the top 50 pathways for each disease and counted how many of these included GOBP, GOCC, GOMF, and Reactome pathways. The results are displayed in Figure 4A. In each disease panel, the horizontal bar on the left indicates the number of pathways in each category that made it into the top 50 (shown by the length of the blue bar). The combined blue and orange segments represent the number of pathways that made it into the top 100.

**Figure 4.**
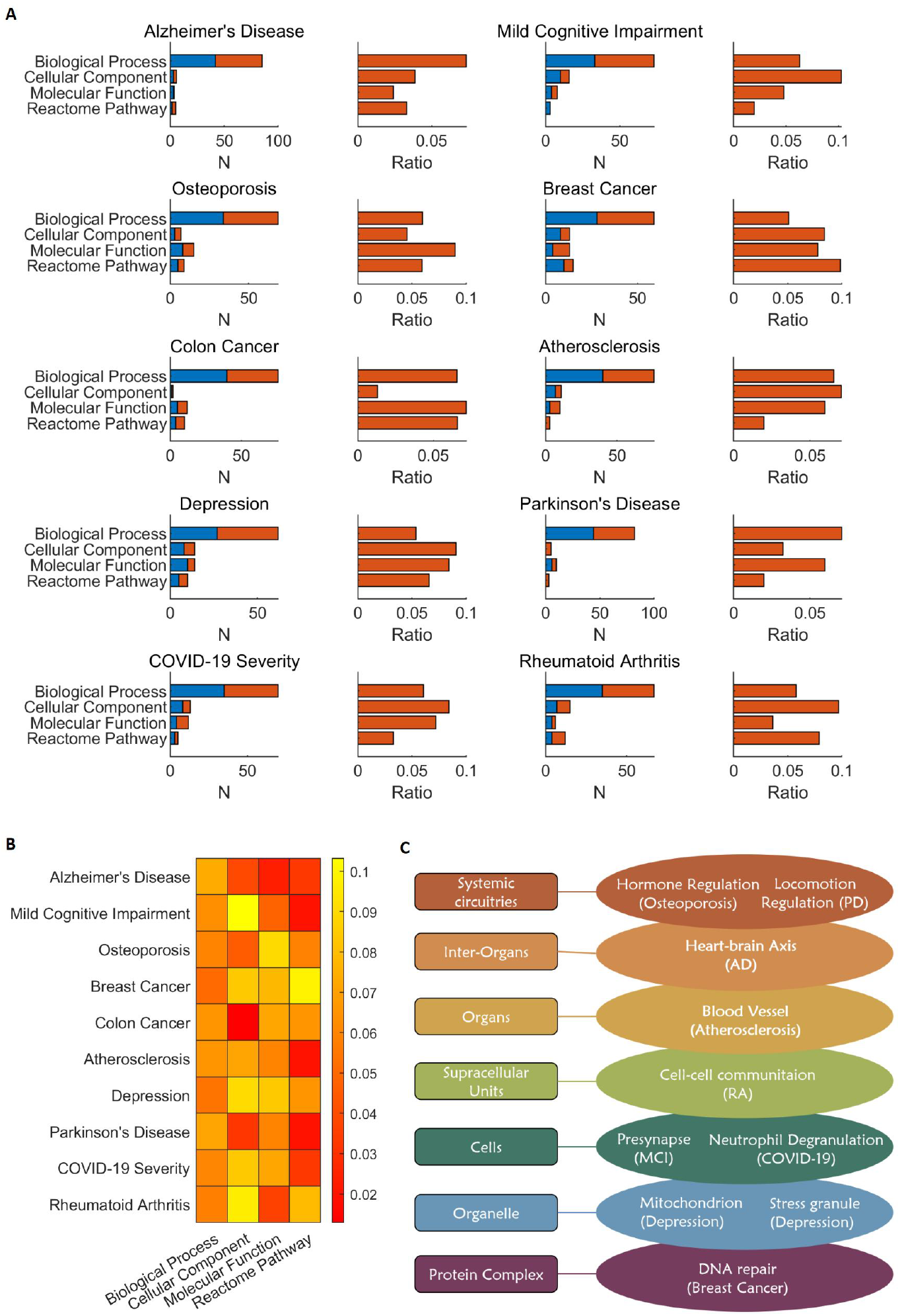
Analysis of Pathway Definitions in Aging-Related Diseases. A. The contribution of pathway terms to the top predictive factors for age-related diseases. B. The relative contribution of pathway terms to diseases. C. Selected pathway terms for creating a multi-scale view of human aging.

Given that the terms defined in GOBP (Gene Ontology Biological Process) are the most numerous, they naturally have a higher likelihood of appearing in the top 100. To account for this, we calculated a ratio for each term type, which is the number of times a term type made it into the top 100 divided by the total number of candidate terms. This ratio reflects the relative likelihood of each term type making it into the top 100 and indicates the relevance of that term type’s aging to the current disease. The ratio value is displayed in the bar chart on the right side of each disease panel.

Among the 10 diseases analyzed, GOBP has the highest absolute number of entries. However, in terms of ratio value, GOBP only ranks first in 2 cases: Alzheimer’s disease and Parkinson’s disease. Interestingly, GOCC, which represents cellular components, ranks first in 5 cases: MCI (mild cognitive impairment), Atherosclerosis, major depressive disorders, COVID-19 severity, and rheumatoid arthritis. GOMF molecular functions rank first in 2 cases (osteoporosis and colon cancer), and the reactome pathway ranks first in one case (breast cancer).

We present the relative ratio value of each category for each disease using a heatmap, as shown in Figure 4B. This visualization clearly demonstrates a strong correlation between the cellular component column and multiple diseases. In omics studies, gene ontology enrichment analysis and pathway enrichment analysis are key bioinformatics tools. Typically, scientists focus on pathway and GOBP analysis, placing less emphasis on the cellular component. Our results suggest that the correlation between cellular components and aging-related diseases deserves attention. The review summary of health hallmarks lists spatial compartmentalization (integrity of barriers and containment of local perturbations) as an important factor, emphasizing the role of life structures in maintaining health [24]. We recommend that future disease research consider changes in cellular components and their roles in disease occurrence. The methods we developed here offer valuable tools for exploring the role of cellular component aging in diseases.

When summarizing the important pathways with predictive significance for each of the diseases discussed earlier, we find that these disease-related pathways span various levels of biological systems—from protein complex, to organelle, to cell, to organ, to inter-organ, and even to systemic circuitries, as illustrated in Figure 4C. For example, at the organelle level, mitochondria and stress granules can predict major depressive disorders. At the inter-organ level, the heart-brain axis can predict Alzheimer’s disease. This indicates that the method can generate a multi-scale view of aging factors in most aging-related diseases.

### The Organelle Aging Index Classifies Age-related Diseases into Nuclear-aging and Cytoplasmic-aging Types

Based on our previous analysis, we examined the common traits of various aging-related diseases using the cellular component pathway aging index. We used the odds ratio of the pathway aging index’s disease prediction ability as our calculation unit to create a global analysis diagram. This diagram illustrates the correlation between the cellular component pathway index and ten types of age-related diseases. We calculated an Overall Age-relatedness score (OA score) for each pathway by summing its odds ratio across all diseases. We then generated a heatmap of the cellular component with the highest OA score and its correlation with the ten diseases, as depicted in Figure 5A. Each square’s number indicates the predictability of the corresponding pathway for the related disease. For better visualization, we scaled the odds ratio between 0 and 1, with a color closer to purple indicating higher corresponding odds.

**Figure 5.**
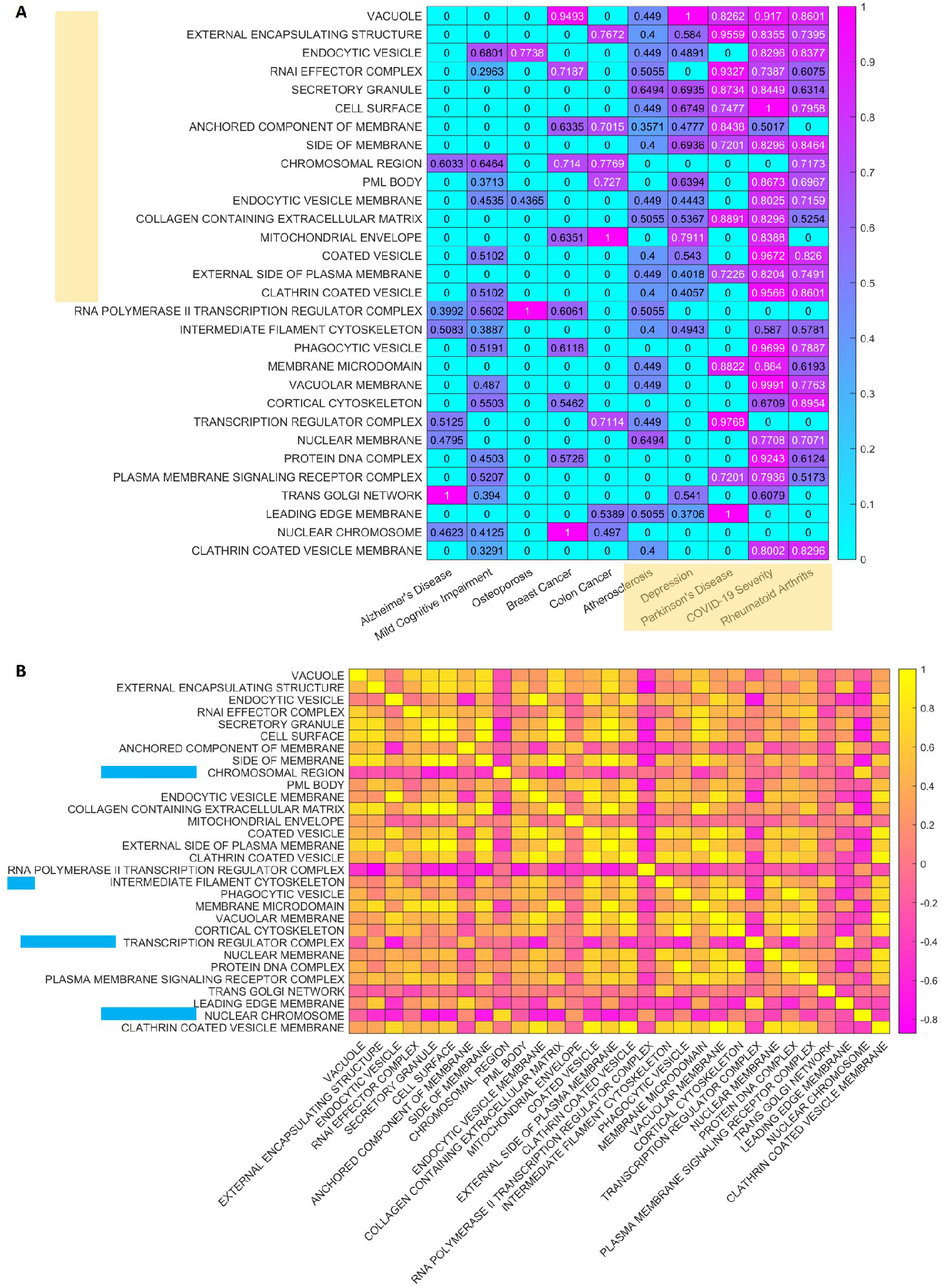
The Cellular Component Aging Index is Linked to Multiple Age-Related Diseases. A. The correlation between the cellular component pathway index and ten types of age-related diseases. B. the correlation between each pathway.

Interestingly, the relationships between many cellular component aging indexes and diseases are not uniformly distributed. The cellular components shown in the figure are mainly associated with the diseases listed on the right half, including Atherosclerosis, Depression, Parkinson’s Disease, COVID-19 severity, and Rheumatoid Arthritis.

We analyzed the correlation between each pathway as shown in Figure 5B. Each pathway, along with its predictive ability for 10 diseases and its odds ratio, is represented as a 10×1 variable. Using Pearson correlation, we calculated the correlation of all pathway combinations and depicted the correlation coefficient rho as a heatmap. Squares closer to yellow indicate correlation coefficients near 1, while rose-colored squares indicate coefficients closer to -1.

The figure clearly shows that most cellular component aging indices have a positive correlation. However, certain pathways, specifically chromosomal region, RNA polymerase II transcription regulator complex, transcription regulator complex, and nuclear chromosome, exhibit a negative correlation.

Considering that in 10 age-related diseases, the cellular component aging index forms two distinct groups, sometimes showing a negative correlation, we selected three representatives from each group. We analyzed their relationship with the 10 diseases, resulting in a clustergram shown in Figure 6A. Interestingly, these six representative cellular component aging indices distinctly categorize all 10 diseases into two primary groups, suggesting the existence of two types of age-related diseases.

**Figure 6.**
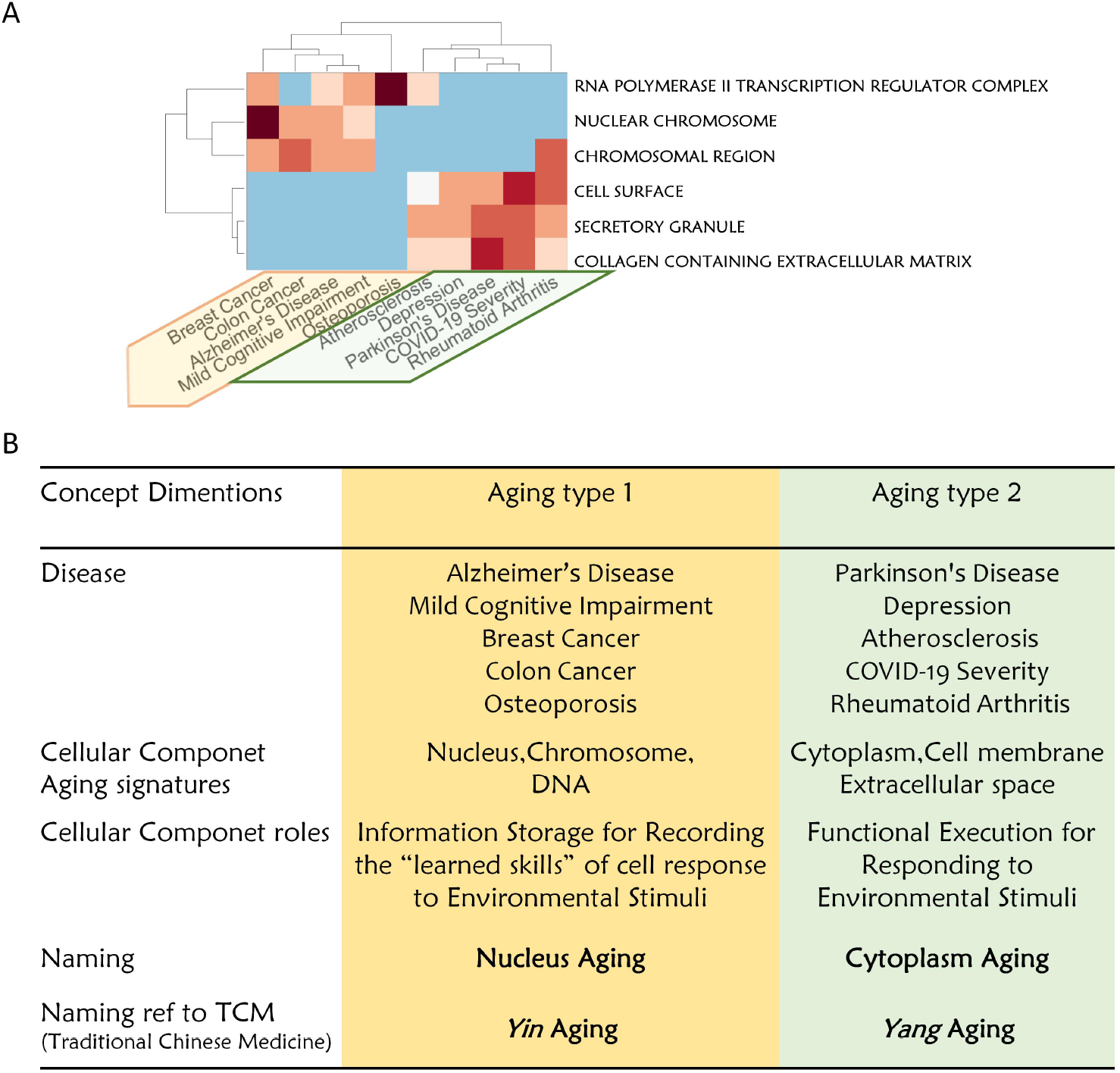
The Cellular Component Aging Index Categorizes 10 Age-related Diseases Into Nucleus Aging and Cytoplasm Aging. A. The clustergram clearly groups these 10 age-related diseases into two clusters. B. A comparative analysis of the two aging types.

The first category includes Alzheimer’s Disease, Mild Cognitive Impairment, Breast Cancer, Colon Cancer, and Osteoporosis. These diseases are linked to aging processes involving the nucleus, chromosome, and DNA. The second category encompasses Parkinson’s Disease, Depression, Atherosclerosis, COVID-19 Severity, and Rheumatoid Arthritis, which are associated with aging processes related to the cytoplasm, cell membrane, and extracellular space.

Interestingly, Alzheimer’s and Parkinson’s diseases, both neurodegenerative conditions, fall into two distinct categories concerning organelle or protein complex aging. We propose naming these two types of aging “nucleus aging” and “cytoplasm aging,” or alternatively, “yin aging” and “yang aging.” We suggest that these different cellular component aging patterns lead to different outcomes. Nucleus aging results in the dysregulation of genetic and epigenetic information storage mechanisms, while cytoplasmic aging represents a decrease in the cell’s functional execution capabilities (Fig 6B).

## Discussion

The aging clock is a dynamic and rapidly advancing research area. Traditionally, it assigns a single value to the entire human body [3]. A recent study introduced ‘10 organ clocks,’ connecting plasma proteome aging signatures with more specific disease predictions for organs [11]. Our paper explores aging in greater detail, calculating it from the organ level down to the cellular, organelle, and even protein complex levels. Expanding from 10 organs to over 3000 pathways presents significant opportunities for biomarker discovery and identifying therapeutic targets.

Aging biomarkers play a vital role in assessing age-related changes, monitoring physiological aging, predicting pathological transitions, and evaluating longevity interventions. Despite the development of numerous aging biomarkers, challenges remain in terms of terminology, validation, and clinical application. Key papers have summarized this field [1, 2], and consortium efforts have been initiated [25]. We believe that pathway aging models, which are data-driven and have multi-scale characteristics, can significantly contribute to this discovery process.

Regarding the scientific mechanism of aging, the “hallmarks of aging” concept represents our current understanding [19]. This article provides a data-driven perspective on these hallmarks, beginning with patterns in 10 age-related diseases. Pathways that predict these diseases align with the hallmarks: DNA repair (genomic instability) predicts Alzheimer’s and breast cancer; protein folding (loss of proteostasis) predicts colon cancer; mitochondrion organization (mitochondrial dysfunction) predicts major depressive disorders; and cell-cell communication (altered intercellular communication) predicts rheumatoid arthritis. Our findings also align with the “hallmarks of health” [24]. We discovered links between membrane aging and disease, which correspond to spatial compartmentalization in the hallmarks of health. Classifying 10 age-related diseases into nucleus and cytoplasm aging provides new tools for understanding diseases at the organelle and molecular levels.

In 2015, the WHO introduced intrinsic capacity (IC) to define healthy aging, covering physical and mental abilities and their interaction with the environment. IC measures the capabilities of aging-relevant biological systems rather than deficits and guides clinical and public health actions. Assessing IC helps understand functional changes and tailor interventions by testing five domains: locomotor, sensory, vitality, psychological, and cognitive. However, scientists sometimes find standardizing IC measures challenging due to heterogeneous tests and reliance on subjective questionnaires and evaluations [26].

We propose that pathway aging models may offer new options in this field. We refer to this method as ‘capomics’ analysis, where capomics stands for the ‘omics of capability’. For any pathway, we calculate the Pathway Aging Index (PA index) and normalize it between 0 and 1. Then, we define a capability index for each pathway (Cap index) using the formula: Cap index = 1 - PA. This enables us to predict phenotypes using the capability index of thousands of pathways.

This method is data-driven and multi-scale, covering everything from protein complexes to inter-organ interactions. It has the potential to uncover causal factors for phenotype changes in diseases. For instance, this study demonstrates that the aging index of heat response pathways can predict Parkinson’s disease, linking it to body temperature control issues, a common symptom of Parkinson’s.

Another advantage of capomics analysis is its convenience and ease of standardization. Unlike comprehensive IC assessments that require hours of collaboration and subjective scales, capomics analysis only needs a single blood or saliva sample, saving time for both clinicians and patients.

The limitations of this article include the study of a relatively small number of age-related diseases. Testing with a broader range of disease types and a larger number of clinical samples is necessary. Additionally, it would be beneficial to investigate whether the method proposed in this article can accurately reflect drug treatment efficacy and their mechanisms of action. This includes tracking changes in aging intervention and the disease recovery process.

## Methods

### Generic Pathway Aging Models

As depicted in Fig.1, we downloaded pathway definitions from The Molecular Signatures Database (MSigDB) (http://www.gsea-msigdb.org/gsea/msigdb/index.jsp). These include gene set definitions from various sources such as Gene Ontology (GO), KEGG pathway, REACTOME pathways, Pathway Interaction Database (PID), WikiPathways (WP), BIOCARTA pathways, and the Hallmark gene set. Notably, Gene Ontology is further divided into three functional categories: Biological Process (GOBP), Molecular Function (GOMF), and Cellular Component (GOCC). For clarity, we use the term “pathway” to refer to the cellular functions performed by all these Gene sets.

We divide all genes within the pathway into signature genes positively associated with aging (Pos-genes) and signature genes negatively associated with aging (Neg-genes). Specifically, we first obtained genome-wide DNA methylation data from GEO data GSE40279. This dataset uses the Illumina Infinium 450k Human DNA methylation Beadchip on whole blood samples of 656 individuals [6]. We compute the average value of methylation sites in each gene’s promoter region as its methylation feature value. Then, we calculate the Pearson correlation between each gene’s methylation feature value and the chronological age of the subjects.

The aging of each pathway is indicated by the statistical difference between the DNA methylation values of Pos-genes and Neg-genes. A T-test is conducted between the DNA methylation values of Pos-genes and Neg-genes for each DNA methylation profile. The resulting T-statistics are used as the pathway aging index (PA index). Specifically, **x** represents DNA methylation values of Pos-genes, and **y** represents DNA methylation values of Neg-genes. The two-class t-test of gene scores (DNA methylation score) between set **x** and set **y** is calculated by T-statistics:

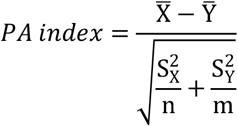

Here, 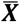 is the mean of gene scores of gene set ***X***, 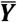 is the mean of gene scores of gene set ***Y***. 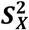 is the variance of gene set X scores, 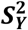 is the variance of gene set ***Y*** scores, ***n*** is the number of genes in set ***X***, and ***m*** is the number of genes in set ***Y***.

### Genome-wide DNA Methylation Data of Ten Age-Related Diseases

We interrogated 10 age-related diseases using genome-wide DNA methylation profiling of clinical cohorts. The data was downloaded from the GEO database. Unless specified otherwise, these data are derived from whole blood samples, using the Illumina 450k/850k microarray platform. The total number of clinical samples is 3263. The data examined in this study encompasses several conditions, including Alzheimer’s Disease [27] (GSE153712, N=632), Mild Cognitive Impairment [27] (GSE153712, N=565), Parkinson’s Disease [28] (GSE111223, N=259, Saliva), Osteoporosis [29](GSE99624, N=48), Breast Cancer (GSE51032, N=659), Colon Cancer (GSE51032, N=590), Atherosclerosis [30] (GSE46394, N=49), Depression [31](GSE113725, N=194), COVID-19 Severity [32](GSE179325, N=473), and Rheumatoid Arthritis [33] (N=689).

Most of the clinical data provided includes both a disease group and a healthy control group, with the exception of the COVID-19 severity data. This particular dataset includes 473 COVID-19 positive individuals. Patients are classified based on the WHO clinical ordinal scale: the mild group (PCR positive individuals, either ambulatory or hospitalized with mild symptoms, on scales 1-4), and the severe group (PCR positive individuals, hospitalized with severe symptoms or who have died, on scales 5-8) [32]. The aim of this analysis is to determine whether the pathway aging index can predict the severity of a patient’s condition, specifically predicting if it will be severe or mild.

The data for breast cancer and colon cancer comes from prospective research involving 845 participants. They were analyzed using a 450k methylation microarray (GSE51032). According to the dataset description on the GEO website, at the time of the last follow-up, 424 participants remained cancer-free. Out of the remaining participants, 235 developed primary breast cancer and 166 developed primary colorectal cancer. Therefore, in the breast cancer dataset, the class label differentiates between the disease group (235) and the control group (424). Similarly, in the colon cancer dataset, the class label differentiates between the disease group (166) and the control group (424).

### Disease Specific Pathway Aging Models

For each disease dataset, we calculate and obtain a set of genes significantly related to the disease phenotype. More specifically, the average value of all methylation sites in each gene promoter region serves as the gene’s methylation value. We use the t-test to calculate the significance of the difference between each gene’s methylation value in the disease group versus the control group. The significantly different genes serve as the disease-related gene set.

For a specific pathway, the aforementioned Pos-genes, i.e., genes positively correlated with age, calculate the intersection with the current disease positively correlated gene set to form disease specific Pos-genes, abbreviated as dsPos-genes. Similarly, genes negatively correlated with age in the pathway, i.e., Neg-genes, calculate the intersection with the current disease negatively correlated gene set to form disease specific Neg-genes, abbreviated as dsNeg-genes.

The aging of each pathway can be indicated by the statistical difference between the DNA methylation values of disease-specific Pos-genes and Neg-genes. This is similar to the pathway aging index calculation described previously in the method part of generic pathway aging models. Specifically, a T-test is conducted between the DNA methylation values of dsPos-genes and dsNeg-genes for each DNA methylation profile. The resulting T-statistics are used as the pathway aging index (PA index). The PA index is then rescaled to a range between 0 and 1.

### Using Pathway Aging Index to Predict Disease

After the above calculations, the methylation dataset for each disease is transformed from a gene×samples matrix to a Pa-index × samples matrix. For each pathway, we use a T-test to determine the difference in the pathway aging index between the disease and control groups. We then use the odds ratio to assess each pathway index’s predictive power for the disease.

The calculation process for the odds ratio is as follows: We calculate the average pathway aging index across all samples in the dataset. Samples with values higher than the average are classified as the PA-High group, while those with lower values are classified as the PA-Low group. Finally, we use the Fisher exact test to determine the difference in disease probability between the PA-High and PA-Low groups, and we obtain the odds ratio value.

## Conflicts of Interest

DeepoMe is a commercial organization developing explainable artificial intelligence (XAI) solutions for health tracking, intervention and drug repurposing in aging related diseases.

## Acknowledgement

This paper has not received any external funding.

## Notes

### Summary of Updates

We added one main figure and improved the results and discussion sections.

